# Human noise blindness drives suboptimal cognitive inference

**DOI:** 10.1101/268045

**Authors:** Santiago Herce Castañón, Dan Bang, Rani Moran, Jacqueline Ding, Tobias Egner, Christopher Summerfield

## Abstract

Humans typically make near-optimal sensorimotor judgments but show systematic biases when making more cognitive judgments. Here we test the hypothesis that, while humans are sensitive to the noise present during early sensory processing, the “optimality gap” arises because they are blind to noise introduced by later cognitive integration of variable or discordant pieces of information. In six psychophysical experiments, human observers judged the average orientation of an array of contrast gratings. We varied the stimulus contrast (encoding noise) and orientation variability (integration noise) of the array. Participants adapted near-optimally to changes in encoding noise, but, under increased integration noise, displayed a range of suboptimal behaviours: they ignored stimulus base rates, reported excessive confidence in their choices, and refrained from opting out of objectively difficult trials. These overconfident behaviours were captured by a Bayesian model which is blind to integration noise. Our study provides a computationally grounded explanation of suboptimal cognitive inferences.

---

The question of whether humans make optimal choices has received considerable attention in the neural, cognitive and behavioural sciences. On one hand, the general consensus in sensory psychophysics and sensorimotor neuroscience is that choices are near-optimal. For example, humans have been shown to combine different sources of stimulus information in a statistically near-optimal manner, weighting each source by its reliability (Ernst & Banks, 2002; Knill, Kersten, & Yuille, 1996; Körding & Wolpert, 2006; Ma, Beck, Latham, & Pouget, 2006; Mamassian, Landy, & Maloney, 2002; Trommershäuser, Maloney, & Landy, 2008). Humans have also been shown to near-optimally utilise knowledge about stimulus base rates to resolve stimulus ambiguity (Kersten, Mamassian, & Yuille, 2004; Körding & Wolpert, 2004; O’Reilly, Jbabdi, Rushworth, & Behrens, 2013; Sun & Perona, 1998; Vilares, Howard, Fernandes, Gottfried, & Kording, 2012).

On the other hand, psychologists and behavioural economists, studying more cognitive judgments, have argued that human choices are suboptimal (Tversky & Kahneman, 1974). For example, when required to guess a person’s occupation, humans neglect the base rate of different professions and solely rely on the person’s description provided by the experimenter. Such suboptimality has been attributed to insufficient past experience (Hertwig & Erev, 2009), limited stakes in laboratory settings (Levitt & List, 2007), the format in which problems are posed (Jarvstad, Hahn, Rushton, & Warren, 2013), distortions in representations of values and probabilities (Ackermann & Landy, 2014), and/or a reluctance to employ costly cognitive resources (Gershman, Horvitz, & Tenenbaum, 2015; Kahneman, 2011). However, an account of human decision-making that can explain both perceptual optimality and cognitive suboptimality has yet to emerge (Summerfield & Tsetsos, 2015).

Here we propose that resolving this apparent paradox requires recognizing that perceptual and cognitive choices often are corrupted by different sources of noise. More specifically, choices in perceptual and cognitive tasks tend to be corrupted by noise which arises at different stages of the information processing leading up to a choice (Faisal & Wolpert, 2009; Hunt, 2014; Juslin & Olsson, 1997; Ma & Jazayeri, 2014). In perceptual tasks, experimenters typically manipulate noise arising before or during sensory encoding. For example, they may vary the contrast of a grating, or the net motion energy in a random dot kinematogram, which affects the signal-to-noise ratio of the encoded stimulus and in turn the sensory percept. Conversely, in cognitive tasks, which often involve written materials or clearly perceptible stimuli, experimenters typically seek to manipulate noise arising after stimulus encoding. For example, they may vary the discrepancy between different pieces of information bearing on a choice, such as the relative costs and benefits of a consumer product (Kahneman, 2011). These types of judgment are difficult because they require integration of multiple, sometimes highly discordant, pieces of information within a limited-capacity system (Botvinick, Braver, Barch, Carter, & Cohen, 2001; Eriksen & Eriksen, 1974; MacLeod, 1991).

Here we test the hypothesis that, while humans are sensitive to noise arising during early sensory encoding, they are blind to the additional noise introduced by their own cognitive system when integrating variable or discordant pieces of information. We tested this hypothesis using a novel psychophysical paradigm which separates, within a single task, these two types of noise. In particular, observers were asked to categorise the average tilt of an array of gratings. We manipulated encoding noise (i.e. the perceptual difficulty of encoding an individual piece of information) by changing the contrast of the array of gratings, with decisions being harder for low-contrast arrays. Second, we manipulated integration noise (i.e. the cognitive difficulty of integrating multiple pieces of information) by changing the variability of the orientations of individual gratings, with decisions being harder for high-variability arrays. Manipulating these different sources of noise within a single task allows us to rule out previous explanations of the optimality gap which hinge on task differences. To pre-empt our results, we show that, while observers adapt near-optimally to increases in encoding noise, they fail to adapt to increases in integration noise. We argue that such “noise blindness” is a major driver of suboptimal inference and may explain the gap in optimality between perceptual and cognitive judgments.

## Results

### Experimental dissociation of encoding noise and integration noise

All six experiments were based on the same psychophysical task (see Methods). On each trial, participants were presented with eight tilted gratings organized in a circular array. Participants were required to categorise the average orientation of the array as oriented clockwise (CW) or counter-clockwise (CCW) from the horizontal axis (**Fig. 1A-B**). After having made a response, participants received categorical feedback about choice accuracy, before continuing to the next trial. We manipulated two features of the stimulus array to dissociate encoding noise and integration noise: the *contrast* of the gratings (root mean square contrast, *rmc*: {0.15, 0.6}), which affects encoding noise, and the *variability* of the gratings’ orientations (standard deviation of orientations, *std*: {0°, 4°, 10°}), which affects integration noise. The distribution of average orientations was identical for all experimental conditions.

**Fig. 1.**
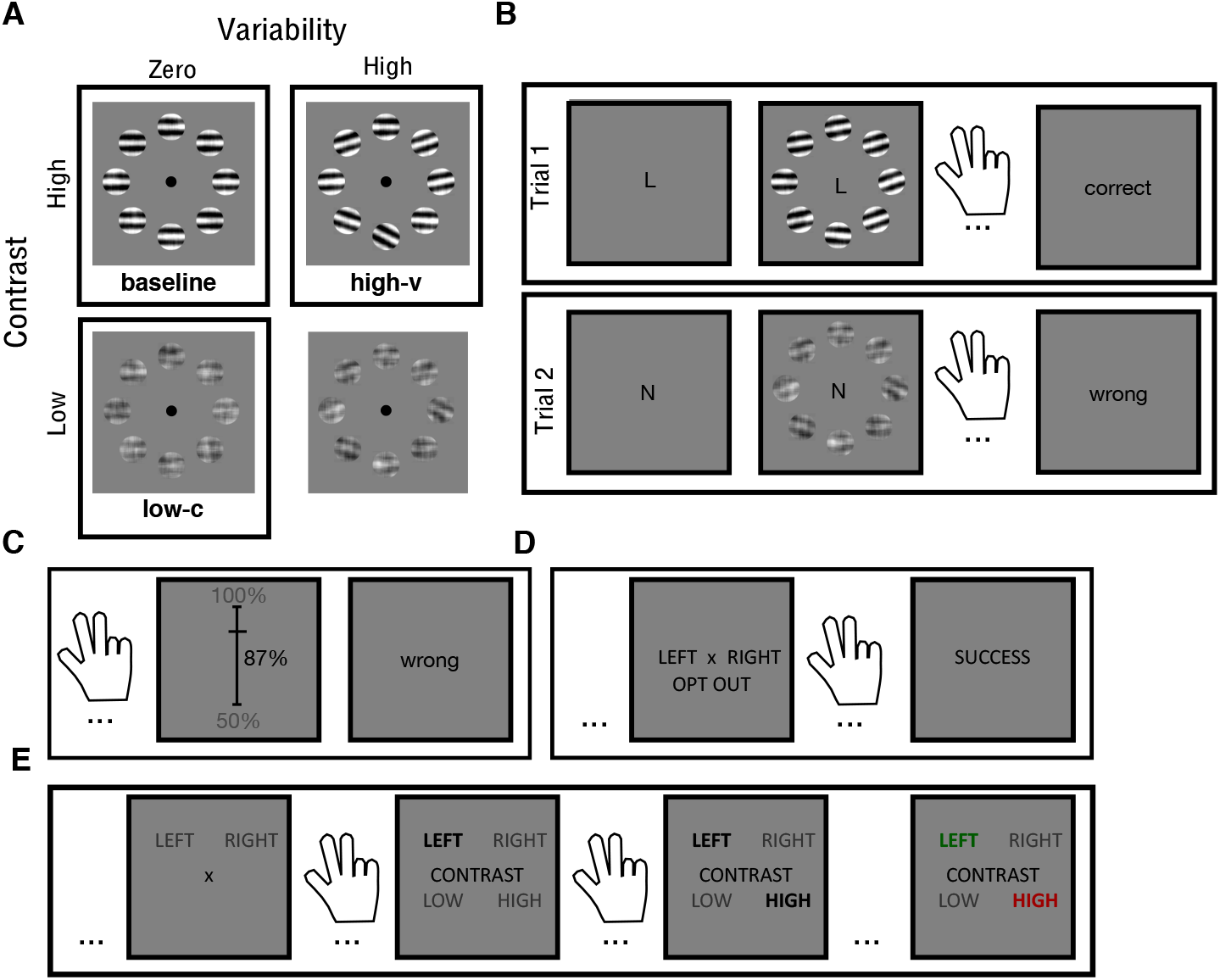
Experimental paradigm. (**A**) We manipulated the stimulus contrast and the orientation variability of an array of eight gratings in a factorial manner. Here we highlight the three critical conditions. (**B**) Participants categorized the average orientation of the array as clockwise (CW, “left”) or counter-clockwise (CCW, “right”) relative to horizontal. A cue, which was shown at the start of each trial and remained on the screen until a response had been made, indicated the prior probability of occurrence of each stimulus category (L: 25% CW, 75% CCW; N: 50% CW, 50% CCW; R: 75% CW, 25% CCW). Participants received categorical feedback about choice accuracy, before continuing to the next trial. Feedback was based on the average orientation of the displayed array. (**C**) In Experiment 2, after having made a choice, participants estimated the probability that the choice was correct by sliding a marker along a scale (50% to 100% in increments of 1%). (**D**) In Experiment 3, participants could opt out of making a choice and receive “correct” feedback with a 75% probability. (**E**) In Experiment 4, after having made a choice, participants were required to categorise (low versus high) either the contrast or the variability of the stimulus array. Here we show a “contrast” trial.

In Experiments 1 (*n* = 20) and 2 (*n* = 20), we assessed the effects of contrast and variability on choice accuracy and evaluated participants’ awareness of these effects. In both experiments, at the beginning of a trial, we provided a “prior” cue which, on half of the trials, signalled the correct stimulus category with 75% probability (henceforth “biased” trials), and, on the other half of trials, provided no information about the stimulus category (henceforth “neutral” trials) (**Fig. 1B**). The neutral trials provided us with a baseline measure of participants’ choice accuracy in the different conditions of our factorial design, and the biased trials allowed us to assess the degree to which – if at all – participants compensated for reduced choice accuracy in a given experimental condition by relying more on the prior cue. In Experiment 2, to provide additional insight into participants’ awareness of their own performance, we also asked participants to report their confidence in the choice (i.e. the probability that a choice is correct; **Fig. 1C**).

### Matched performance _ for different levels of encoding and integration noise

We first used the neutral trials to benchmark the effects of contrast and variability on choice accuracy. As intended, choice accuracy decreased with lower contrast (Exp1: *F*(1,19) = 15.54, *p* < .001; Exp2: *F*(1,19) = 41.08, *p* < .001; collapsed: *F*(1,39) = 49.3, *p* < .001) and with higher variability (Exp1: *F*(1.3,24.7) = 8.51, *p* < .001; Exp2: *F*(1.6,32.2) = 26.0, *p* < .001; collapsed: *F*(1.4,57.3) = 30.61, *p* < .001). Our factorial design contained three critical conditions which allowed us to compare participants’ behaviour under distinct sources of noise: (i) “baseline”, (ii) “low-c” and (iii) “high-v”. In the baseline condition, the total amount of noise is lowest (high contrast, .6; zero variability, 0°). In the low-c condition (low contrast, .15; zero variability, 0°), encoding noise is high but integration noise is low. Conversely, in the high-v condition, integration noise is high but encoding noise is low (high contrast, 0.6; high variability, 10°). As expected, choice accuracy was reduced both in the low-c and in the high-v conditions (about 12%) compared to the baseline condition (baseline>low-c: *t*(39) = 9.24, *p* < .001; baseline>high-v: *t*(39) = 9.70, *p* < .001; **Fig. 2A**). Critically, choice accuracy was at statistically similar levels in the low-c and the high-v conditions (Exp1, high-v>low-c: *t*(19) = 0.36, *p* > *0.7;* Exp2, high-v>low-c: *t*(19) = 0.11, *p* > 0.9; collapsed, high-v>low-c: *t*(39) = 0.34, *p* > 0.7; **Fig. 2A**). Overall, the results show that we successfully manipulated noise at different stages of information processing.

**Fig. 2.**
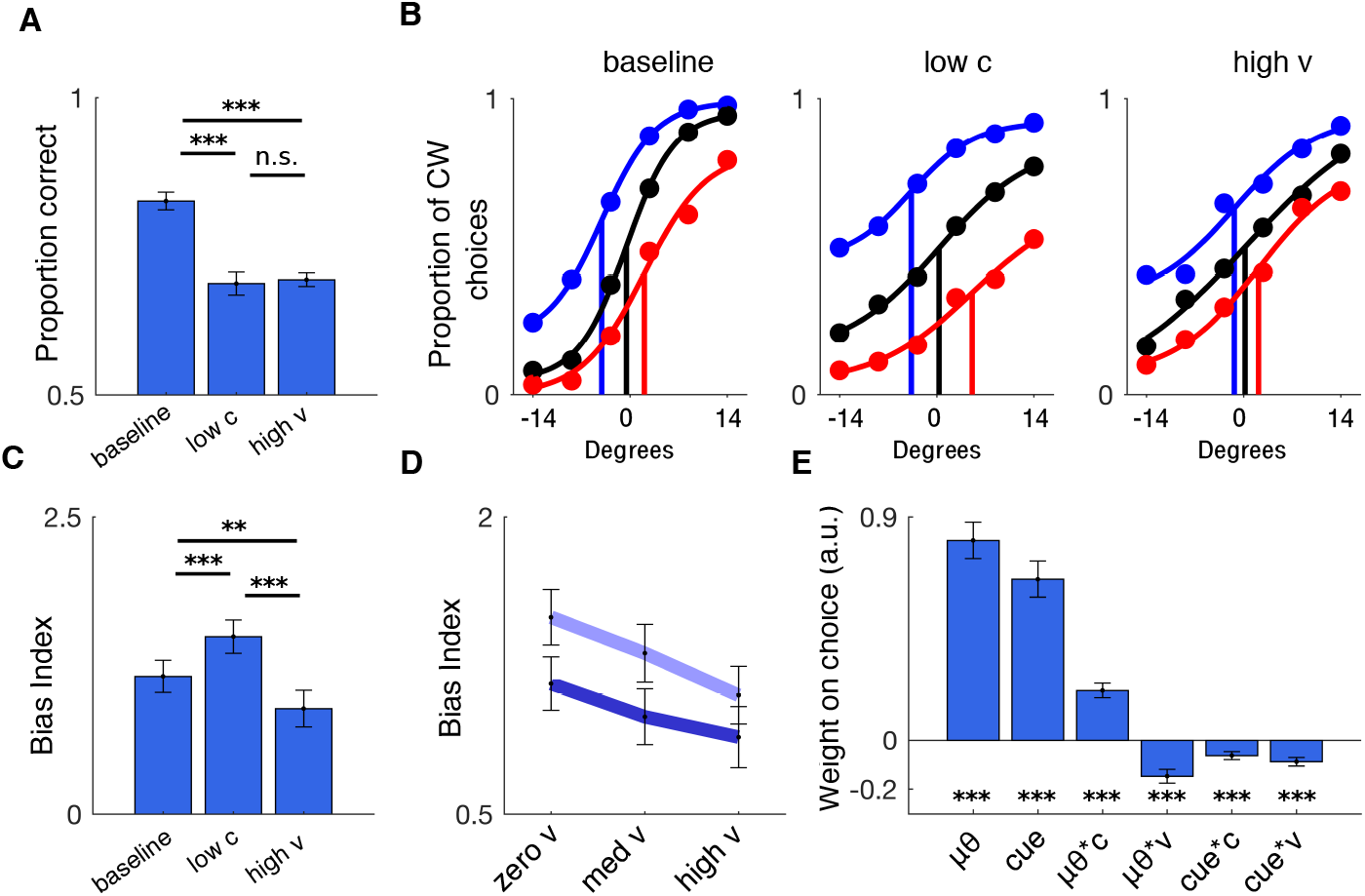
Effects of contrast and variability on choice behaviour. (**A**) Choice accuracy for the baseline, reduced contrast (low-c) and increased variability (high-v) conditions. (**B**) Psychometric curves are shallower in the low-c and high-v conditions compared to baseline. The x-axis indicates average orientation relative to horizontal, with negative and positive values for CCW and CW, respectively. Choices shift towards the cued category on biased trials (blue: 75% CW; red: 25% CW) compared to neutral trials (black) but least so in the high-v condition. Vertical lines mark the inflection points of psychometric functions fitted to the average data. Psychometric curves were created for illustration. (**C**) Bias index, a measure of cue usage, is higher in the low-c condition but lower in the high-v condition compared to baseline. (**D**) Factorial analysis of the effects of contrast and variability on the bias index shows an increase with contrast (dark blue: high contrast; pale blue: low contrast) but a decrease with variability. (**E**) Trial-by-trial influence of prior cue on choices measured using logistic regression. c: contrast; v: variability; μθ: signed mean orientation; cue: signed prior cue. (**A-E**) Data is represented as group mean ± SEM. \**p* < .05, \*\**p* < .01, \*\*\**p* < .001. For panel A, only neutral trials were used. For panel B and E, both neutral and biased trials were used. For panels C and D, only biased trials were used.

### Do people utilise the prior cue to compensate for increased errors?

We next leveraged the biased trials to assess the degree to which participants adapted to the changes in choice accuracy induced by our factorial design. Given the above results, we would expect participants to rely more on the prior cue in the low-c and the high-v condition than in the baseline condition. To test this prediction, we applied Signal Detection Theory (Macmillan & Creelman, 2004; Stanislaw & Todorov, 1999) to quantify the degree to which participants shifted their decision criterion in accordance with the prior cue (see Methods). Briefly, we constructed a “bias index” computed as the difference in the decision criteria between the condition in which the prior cue was “clockwise” and the condition in which the prior cue was “counter-clockwise”. The higher the bias index, the higher the influence of the prior cue on choice. As expected under an ideal observer framework, participants used the prior cue more in the low-c than in the baseline condition (*t*(39) = 4.89, *p* < .001; **Fig. 2C**). However, contrary to an ideal observer framework, participants used the prior cue less in the high-v than in the baseline condition (*t*(39) = 2.85, *p* < .01; **Fig. 2C**). This pattern is clear from the psychometric curves constructed separately for each condition shown in **Fig. 2B** (compare inflection points).

In line with these results, a full factorial analysis of the bias index identified a positive main effect of contrast (*F*(1,39) = 24.02, *p* < .001) and a negative main effect of variability (*F*(1.9,37.1) = 9.9, *p* < .001; **Fig. 2D**). Finally, including both neutral and biased trials, we used trial-by-trial logistic regression to investigate how contrast (c) and variability (v) affected the influence of the prior cue and sensory evidence (μθ) on choices (μθ, cue, μθ*c, μθ*v, cue*c, cue*v; **Fig. 2E**). The prior cue had a larger influence on choices on low-contrast compared to high-contrast trials (*t*(39) = 4.05, *p* < .001) and on low-variability compared to high-variability trials (*t*(39) = 5.21, *p* < .001). Taken together, these results show that participants did not adapt to the additional noise arising during integration of discordant pieces of information.

### Are people blind to integration noise?

To test whether participants failed to adapt because they were “blind” to integration noise, we analysed the confidence reports elicited in Experiment 2 (**Fig. 1C**). We implemented a strictly-proper scoring rule such that it was in participants’ best interest (i) to make as many accurate choices as possible and (ii) to estimate the probability that a choice is correct as accurately as possible (Sonnemans & Theo Offerman, 2001). In support of our hypothesis, analysis of the full factorial design showed that, while confidence varied with contrast (*F*(1,19) = 32.97, *p* < . 001), it did not vary with variability (*F*(1.2,22.5) = 0.73, *p* > 0.4). In addition, direct comparison between the low-c and high-v conditions showed that participants were more confident in the high-v condition (*t*(19) = 3.98, *p* < .001; **Fig. 3A**), with participants overestimating their performance (difference between mean confidence and mean accuracy; *t*(19) = 2.66, *p* < .05; **Fig. 3A**). Although participants reported lower confidence in the high-v condition compared to baseline (**Fig. 3A**), this decrease was due to participants utilising response times as a cue to confidence (Zakay & Tuvia, 1998): a trial-by-trial regression analysis showed that confidence decreased with longer response times (RTs) and was unaffected by variability once RTs had been accounted for (v: *t*(19) = 0.38, *p* > 0.7; all other t-values > 4, all *p* < . 001; see **Fig. 3B** and *Response times* in the Supplementary Information). Overall, these results show that participants were overconfident under integration noise, as if they were “blind” to the impact of integration noise on their performance.

**Fig. 3.**
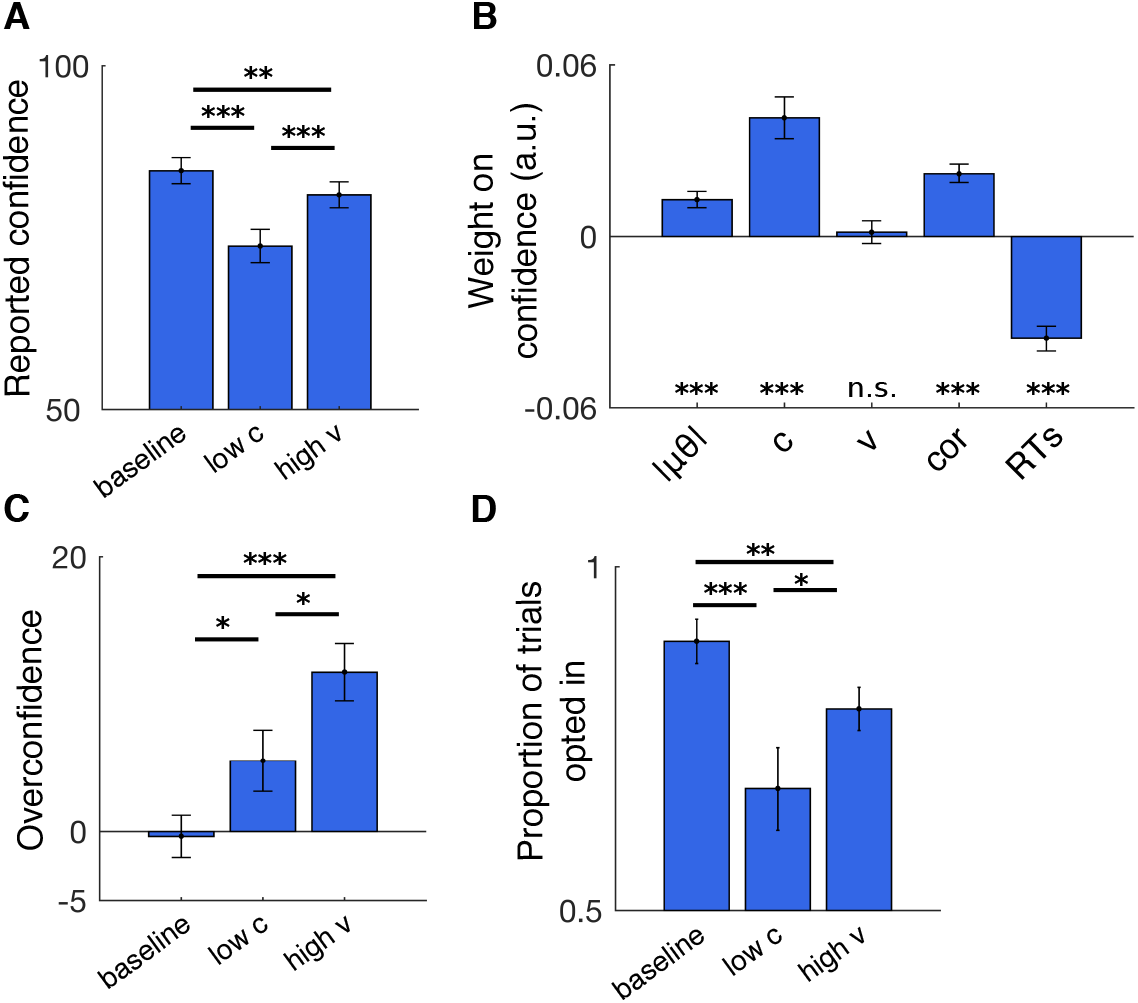
Effects of contrast and variability on explicit and implicit markers of confidence. (**A**) Mean confidence in the baseline, reduced contrast (low-c) and increased variability (high-v) conditions. (**B**) Trial-by-trial confidence is not influenced by variability (v) but is influenced by the deviation of the average orientation from horizontal (|μθ|), contrast (c), choice accuracy (cor) included in order to account for error detection (Yeung & Summerfield, 2012), and response times (RTs). (**C**) Overconfidence, the difference between mean confidence and mean choice accuracy, is highest in the high-v condition. (**D**) Higher probability of making a choice (and thus not opting out) in the high-v condition compared to the low-c condition. (A-D) Data are represented as group mean ± SEM. For panel B, only biased trials were used. For all other panels, only neutral trials were used.

In Experiment 3 (*n* = 18), because explicit confidence reports can be highly idiosyncratic (Aitchison, Bang, Bahrami, & Latham, 2015; Bang et al., 2017), we obtained an implicit, but perhaps more direct measure, of confidence (Hampton, 2001; Kepecs & Mainen, 2012; Kiani & Shadlen, 2009). Specifically, on half of the trials (“optional trials”), we introduced an additional choice option, an opt-out option, which yielded “correct” feedback with a 75% probability. On the other half of trials (“forced trials”), participants had to make an orientation judgment. Under this design, to maximise reward, participants should choose the opt-out option whenever they thought they were less than 75% likely to make a correct choice. Despite matched levels of choice accuracy in the low-c and the high-v conditions (forced trials, *t*(17) = 0.24, *p* > 0.8), participants decided to make an orientation judgment more often on high-v than on low-c trials (optional trials, *t*(17) = 2.32, *p* < .05; **Fig. 3D**), again indicating overconfidence in the face of integration noise. A full factorial analysis verified that the proportion of such opt-in trials varied with contrast (*F*(1,17) = 21.2, *p* < .001) but not with variability (*F*(1.4,23.9) = 3.6, *p* > 0.05). Similarly, a trial-by-trial logistic regression showed that the probability of opting in varied with contrast (*t*(17) = 6.93, *p* < .001) but not with variability (*t*(17) = 1.6, *p* > 0.1), after controlling for other task-relevant factors (e.g., average orientation and RTs). In sum, participants opted out more often when encoding noise was high, but did not do so when integration noise was high, despite making a comparable proportion of errors in the two conditions.

### Computational model of noise blindness

We next compared a set of computational models based on the ideal observer framework to provide a mechanistic explanation for the observed data (see Methods). There are broadly three components to our modelling approach. First, a generative (true) model which describes the task structure and the generation of noisy sensory data. Second, an agent’s internal model of the task structure and how sensory data is generated; the internal model may differ from the generative model. Finally, a Bayesian inference process which involves inverting the internal model in order to estimate the probability of a stimulus category given sensory data and generate a response. This inference process involves marginalising over contrast and variability levels according to a belief distribution over experimental conditions. Optimal behaviour can be said to occur when there is a direct correspondence between the generative model and the agent’s internal model. We evaluated the models both qualitatively (i.e. model predictions for critical experimental conditions) and quantitatively (i.e. BIC scores).

We focus on an “omniscient” model, which has perfect knowledge of the task structure and how sensory data is generated, and two suboptimal models which propose different mechanistic explanations of participants’ lack of sensitivity to the performance cost associated with stimulus variability. The suboptimal models relax the *omniscient* assumptions about an agent’s beliefs about (i) the task structure and/or (ii) the sources of noise in play. See Supplementary Information for details about all models considered.

In our task the average orientation of a stimulus array was sampled from a common distribution of orientations across experimental conditions (**Fig. 4A**). We modelled an agent’s sensory data as a random (noisy) sample from a Gaussian distribution centred on the average orientation of the stimulus array (**Fig. 4B**), with the variance of this distribution determined by both encoding noise and integration noise. We used each participant’s data from the neutral trials to parameterise their levels of encoding noise and integration noise in each experimental condition (see Methods). The fitted noise levels, which are part of the generative model, were the same for all models; thus no additional free parameters were fitted to the data and the models only differed with respect to their assumptions about the internal model used for Bayesian inference.

**Fig. 4.**
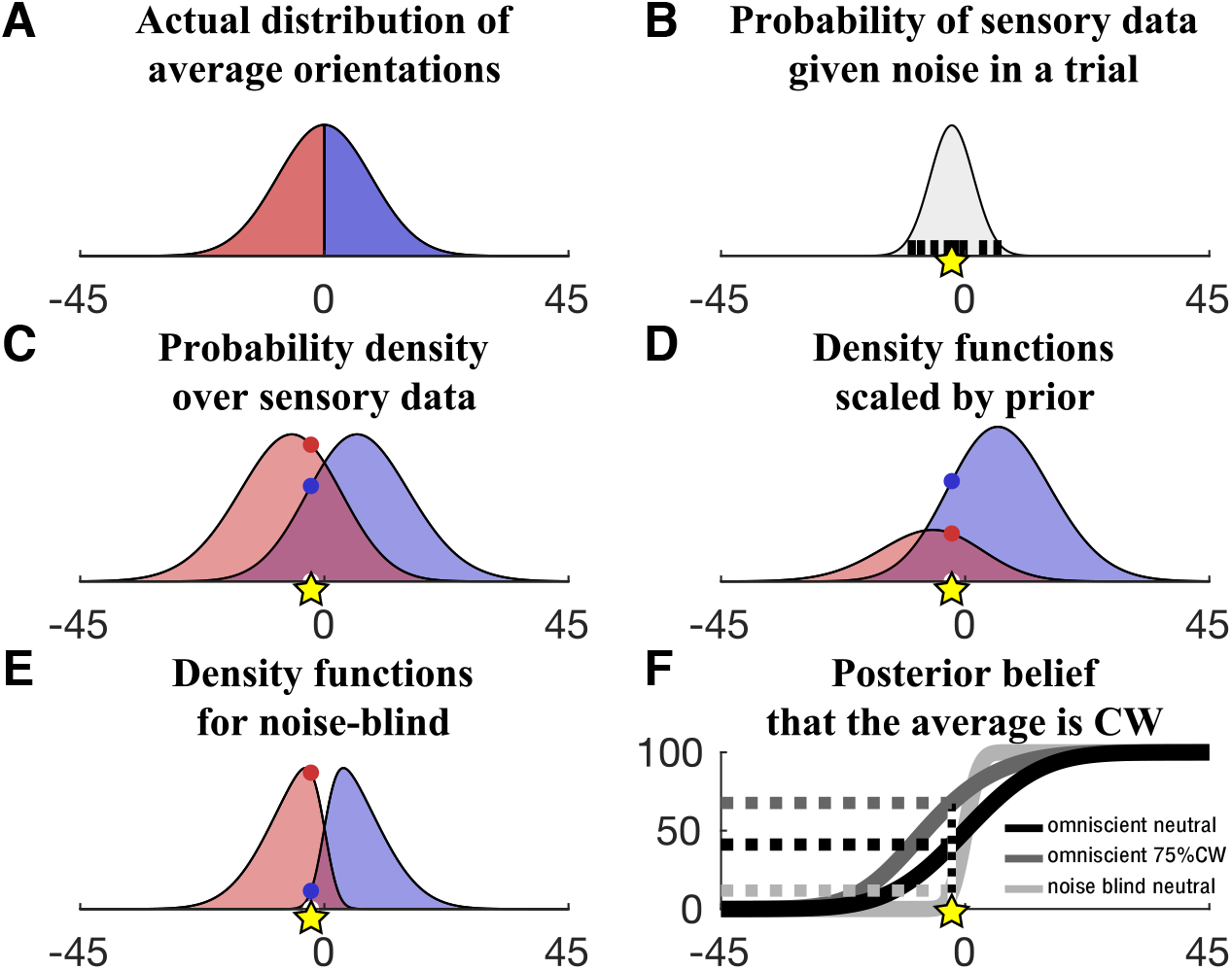
Computational model. (**A**) Distribution of average orientations conditioned on CCW (red) and CW (blue). (**B**) An agent’s sensory data was modelled as a sample from a Gaussian distribution centred on the average orientation of the eight gratings in a stimulus array (black vertical lines), with the variance of this distribution determined by both encoding noise and integration noise. The yellow star marks the sensory data for an example trial. (**C**) An omniscient agent has a pair of category-conditioned probability density functions over sensory data for each experimental condition (i.e. contrast × variability level; here a single condition is shown). The agent uses the relevant pair of density functions to compute the probability of the observed sensory data (yellow star) given each category (red and blue dots). Note that, for an omniscient agent, these density functions match the true probability density over sensory data under the generative model. See an example of a full set of density functions for an experiment in **Fig. S1**. (**D**) Density functions from panel C after scaling by the prior cue (here 75% CW). The sensory data (yellow star) is now more likely to have come from a CW stimulus than a CCW stimulus. (**E**) The noise-blind model only takes into account encoding noise: the density functions therefore overlap less than in panel C and they do not match the true probability density over sensory data. (**F**) Posterior belief that the stimulus is CW as a function of the same sensory data (yellow star) for the examples shown in panels C (black, omniscient model on neutral trials), D (dark grey, omniscient model when prior cue is 75% CW) and E (light grey, noise-blind model on neutral trials). Steeper curves indicate higher confidence; categorisation accuracy (on neutral trials) is the same for all models. The variability-mixer curve would have intermediate slope between that of the omniscient and the noise blind model in conditions of high variability.

The *omniscient* model has, for each experimental condition, a pair of functions that specify the probability density over sensory data given a CW and a CCW stimulus, taking into account both encoding and integration noise. As the model can identify the current condition (e.g., knows with certainty that a trial is drawn from the high-contrast, high-variability condition), it only uses the relevant pair of density functions to compute the probability of the observed sensory data given a CCW and a CW category (**Fig. 4C**). On neutral trials, each category is equally likely, and the agent computes the probability that a stimulus is CW and CCW directly from the density functions. On biased trials, the categories have different prior probabilities, and the agent scales the density functions by the prior probability of each category as indicated by the prior cue (**Fig. 4D**). After having calculated the probability that a stimulus is CW and CCW, the agent can compute a choice (i.e. chose the category with the higher posterior probability) and confidence in this choice (i.e. the probability that the choice is correct)

We now consider two competing explanations of the participants’ lack of sensitivity to the performance cost associated with stimulus variability. First, a *variability-mixer* model which relaxes the assumption that an agent can identify the current variability condition. The model therefore uses a single pair of density functions for all variability conditions (which are a mixture of density functions across variability levels). As a result, compared to the *omniscient* model, the density functions are wider on low-variability trials but narrower on high-variability trials. Second, a *noise-blind* model which relaxes the assumption that the agent is aware of integration noise. As for the variability-mixer model, the noise-blind model uses a single pair of density functions for all variability conditions, but, critically, these density functions do not take into account the additional noise induced by stimulus variability. Because of these differences in the internal model used for Bayesian inference, the models differ in the degree of confidence in a choice for a given sensory data (**Fig. 4F**) and, by extension, the influence of the prior cue on choice on biased trials.

In support of our hypothesis, the noise-blind model provided the best fit to our data. First, the noise-blind model, and not the omniscient model, predicted three key features of participants’ behaviour: (i) overconfidence on high-variability trials within participants (**Fig. S2**), (ii) no correlation between mean accuracy and mean confidence across participants (**Fig. 5A**) and (iii) a diminished influence of the prior cue on high-variability trials, as seen by both the analysis of the bias index (**Fig. 5C**) and the trial-by-trial regression predicting confidence (**Fig. 5D**), where the prior cue has a positive effect on confidence but its effect decreases with high contrast and high variability (in line with noise blindness). In addition, quantitative comparison yielded “very strong evidence” (Kass & Raftery, 1995) for the noise-blind model over the omniscient model, with an average ΔBIC across participants of -32.9 (**Fig. 5B**). Similarly, analyses of the patterns of overconfidence in the critical conditions of our factorial design favoured the noise-blind over the variability-mixer model (**Fig. S2**), and quantitative comparison yielded “very strong evidence” for the noise-blind over the variability-mixer model (ΔBIC = -20.4, **Fig. 5B**). In sum, the modelling indicates that participants neglected integration noise altogether.

**Fig. 5.**
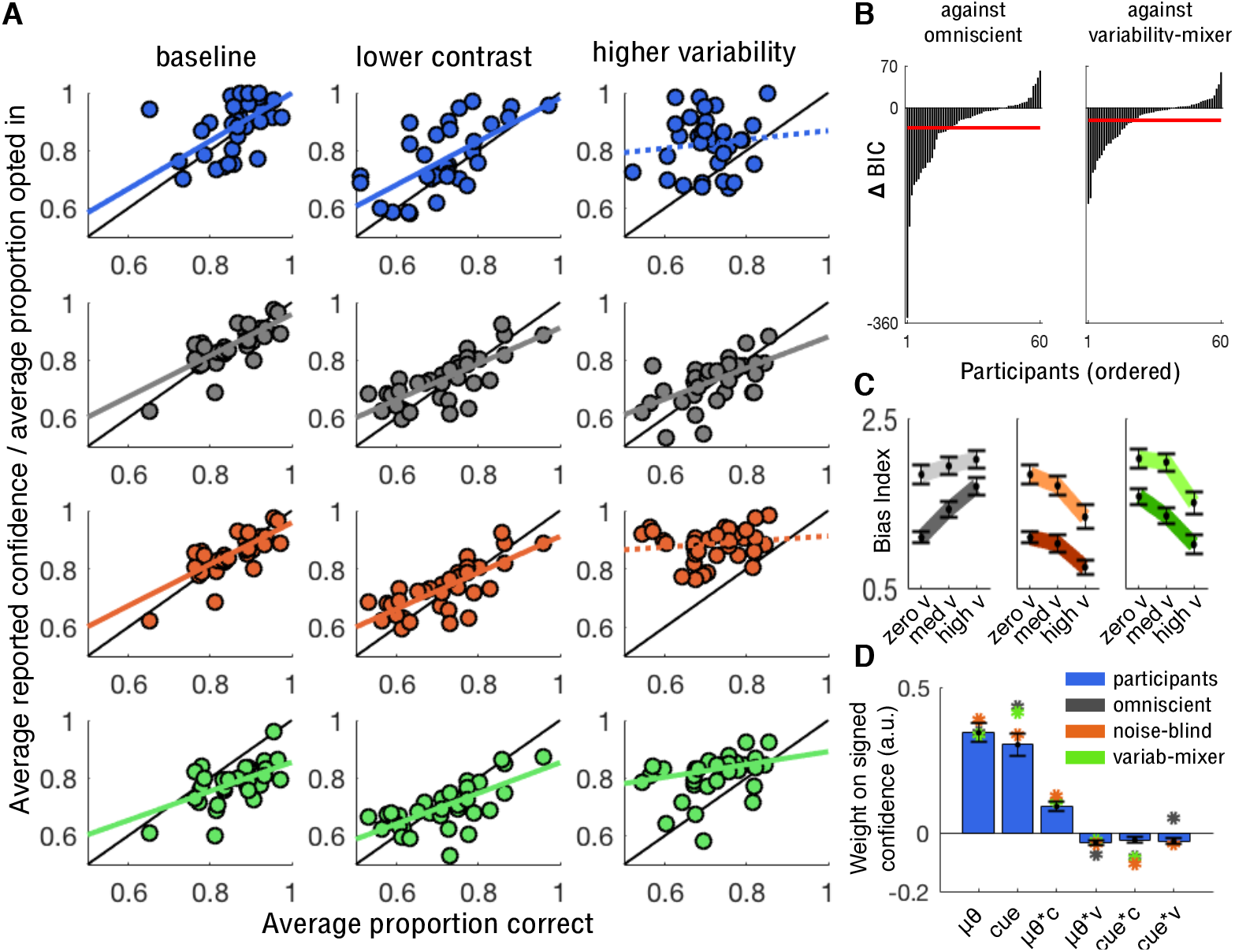
Comparison of model and human behaviour. (**A**) Correspondence between mean accuracy and mean confidence (explicit estimates or proportion of opt-in responses) for participants (blue, data from Exp1-3) and the omniscient (grey), noise-blind (orange) and variability-mixer (green) models in the critical experimental conditions. Coloured lines indicate best-fitting slope of a linear regression analysis: solid for *p* < .05, dotted for *p* > 0.4. (**B**) Model comparison (Exp1-3) suggests strong evidence in favour of the noise-blind model over the omniscient model (left panel, average ΔBIC = -32.9) and also over the variability-mixer model (right panel, ΔBIC = -20.4). (**C**) Omniscient model (left) makes opposite predictions to noise-blind model (middle) and variability-mixer model (right) for the influence of the prior cue on choice (Exp1 - 2) as variability increases (positive versus negative slopes) but similar predictions as contrast decreases (lighter lines above darker ones). Dark colours: high contrast. Light colours: low contrast. (**D**) Trial-by-trial analysis of signed confidence (Exp2; negative for CCW and positive for CW) for participants (blue) and the omniscient (grey), noise-blind (orange) and variability-mixer (green) models. (C-D). Data are represented as group mean ± SEM. For panel A, only neutral and optional trials were used. For panels B and C, only biased trials were used. For panel D, both neutral and biased trials were used. Within-model variability in predictions comes from variability in encoding and integration noise across participants.

### Participants are noise blind and not variability blind

To further rule out the hypothesis that participants were simply unable to discriminate the variability conditions as proposed by the variability-mixer model, we ran Experiment 4 *(n* = 24). After having made a choice, participants were asked to categorise either the contrast of the stimulus array (*rmc*, high: .6 vs. low: .15) or the variability of the stimulus array (*std*, high: 10° vs. low: 0°) (**Fig. 1E**). Again, choice accuracy on neutral trials in the low-c and the high-v conditions was statistically indistinguishable (*t*(23) = 1.16, *p* > 0.2). We reasoned that, if participants had difficulty identifying the variability condition but otherwise aware of integration noise, then they should behave closer to optimal when they correctly identified the variability condition. To test this prediction, we used the biased trials to compare cue usage when the variability condition was correct and incorrectly categorised (75.71% ± 2.26% of the variability-condition judgments were correct). In contrast to the prediction, but in line with our hypothesis, participants showed blindness to integration noise even when they correctly identified the variability condition: participants were more biased on low-c than high-v trials regardless of whether the variability categorisation was correct (*t*(23) = 3.21,*p* < .01) or incorrect (*t*(23) = 4.05,*p* < .001; **Fig. 6A-B**).

**Fig. 6.**
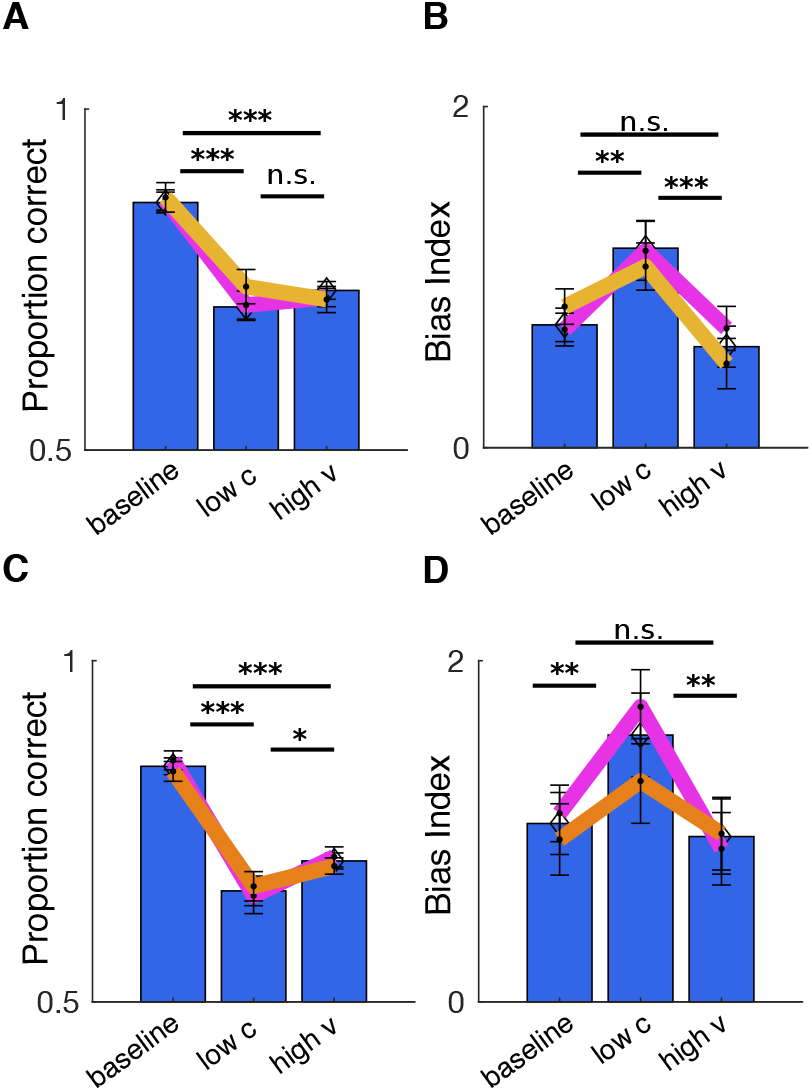
Experimental evidence against variability mixing. (**A**) Choice accuracy for the baseline, reduced contrast (low-c) and increased variability (high-v) conditions for Experiment 4. (**B**) In Experiment 4, the influence of the prior cue is highest when encoding noise is high (low-c) and lowest when integration noise is high (high-v). (**C**) Same as panel A, but for Experiment 5. (**D**) Same as panel B, but for Experiment 5. (A-B) Coloured lines indicate trials where the categorisation of contrast was correct (pink) and where the categorisation of variability was correct (yellow). (C-D) Coloured lines indicate trials where the contrast level was blocked (pink) or when the variability level was blocked (brown). We note that the difference in bias index for the low-c condition between contrast blocking and variability blocking can be explained by a general shift in the bias index according to block difficulty: when contrast is blocked, the low-c condition is accompanied by the hardest condition (the condition with low contrast and high variability), but when variability is blocked, the low-c condition is accompanied by the easiest condition (the condition with high contrast and zero variability). (A-D) Data are represented as group mean ± SEM. For panels A and C, neutral trials were used. For panels B and D, biased trials were used.

In Experiments 1-4, the experimental conditions were interleaved across trials, which may have made it too difficult for participants to separate the different sources of noise in play. To test the generality of our results, we ran Experiment 5 (*n* = 24) in which either the contrast or variability level were kept constant across a block of trials (**Fig. 6C-D**). Even then, and despite receiving trial-by-trial feedback, participants were not, compared to the baseline condition, more influenced by the prior cue when variability was high (biased trials, *t*(23) = 0.32, *p* > 0.7), but they were when contrast was low (biased trials, *t*(23) = 3.31, *p* < .01). In other words, even under blocked conditions participants failed to learn about the performance cost associated with stimulus variability.

### Sequential sampling account of noise blindness

A recent study investigated how stimulus volatility (i.e. changes in evidence intensity across a trial) affected choice and confidence (Zylberberg, Fetsch, & Shadlen, 2016). Participants were found to make faster responses and report higher confidence when stimulus volatility was high. These results were explained by a sequential sampling model which assumes that observers are unaware of stimulus volatility and therefore, unlike an “omniscient” model, adopt a common choice threshold across trial types. In the Supplementary Information, we show, using empirical and computational analyses, that this model cannot explain our results (**Fig. S3**). For example, the model predicts *faster* RTs on high-variability than low-variability trials, a prediction which is at odds with our observation of *slower* RTs on high-variability trials.

### Noise blindness cannot be explained by subsampling

We have proposed that stimulus variability impairs performance because of noise inherent to cognitive integration of variable or discordant pieces of information. An alternative explanation of the performance cost for high stimulus variability is that participants based their responses on a subset of gratings rather than the full array. Under this *subsampling* account, choice accuracy for high-variability stimuli is lower because of a larger mismatch between the average orientation of the full array and the average orientation of the sampled subset. Here we provide several lines of evidence against the subsampling account (see details in Supplementary Information).

We first examined performance under different set-sizes in Experiment 6 (*n* = 20) where the stimulus array was made up of either four or eight gratings (average orientations and orientation variability were equated across set-sizes). We reasoned that, if participants did indeed engage in subsampling, then performance should be higher for four than eight gratings. Because of the matched average tilt in the array, sampling four items would impair performance in the high-v condition for an eight-item array but not for a four-item array. However, we found no effect of set-size on choice accuracy (*F*(1,20) = 0.006, *p* > 0.9; **Fig. S4A**); the effects of contrast (*F*(1,20) = 40.9, *p* < .001) and variability (*F*(1,20) = 30.50, *p* < .001) were comparable to those observed in our previous experiments.

We next simulated performance for eight-grating arrays under a subsampling agent which did not have integration noise but instead sampled a subset of the items (1-8 items, **Fig. S4B**). The observed difference in participants’ performance between the baseline and the high-v conditions could be explained by assuming an agent that sampled about four items out of eight. However, this account – because there is no integration noise – predicts that participants should have similar levels of performance for the baseline and the high-v conditions for four-item arrays, a prediction which is at odds with our data (**Fig. S4A**). If integration noise is introduced, then most, if not all, items would have to be sampled to account for the data.

Finally, we fitted a computational model to participants’ choices in Experiments 1 to 3 (eight-item arrays) in order to directly estimate the number of items sampled by each participant. This modelling approach revealed that the majority participants (42 out of 60) sampled all eight items (Table S2). We note that subsampling, even if an auxiliary cause of integration noise, cannot without further assumptions (e.g. blindness to the performance cost) explain participants’ lack of sensitivity to the performance cost associated with high-variability stimuli.

## Discussion

Here we propose a new explanation for the previously reported gap in optimality between perceptual and cognitive decisions. Using a novel paradigm, we show, within a single task, that humans are sensitive to noise present during sensory encoding, in keeping with previous perceptual studies (Ernst & Banks, 2002; Körding & Wolpert, 2004), but blind to noise arising when having to integrate variable or discordant pieces of information, a typical requirement in cognitive tasks. This noise blindness gave rise to two common signatures of suboptimality often found in cognitive studies: base-rate neglect and overconfidence.

We provided several lines of evidence for our hypothesis. When stimulus variability was high, participants were overconfident, as indicated by cue usage, subjective confidence reports as well as opt-in responses, even though they received trial-by-trial feedback, and even when stimulus variability was salient (Exp1-3), accurately categorised (Exp4) or constant across a block of trials (Exp5). These findings indicate that, while participants were able to track stimulus variability, they simply neglected the performance cost associated with high-variability stimuli. We also ruled out that such noise blindness was due to participants only sampling a subset of a stimulus array (Exp6). The best model of our data assumed that participants sampled all items and were blind to the additional noise inherent to cognitive integration of variable or discordant pieces of information.

An extensive literature has considered the different types of noise which affect human choices (Beck, Ma, Pitkow, Latham, & Pouget, 2012; Hunt, 2014; Juslin & Olsson, 1997). Our classification is partially related to a previous distinction between noise which originates inside the brain, such as intrinsic stochasticity in sensory transduction (Thurstone, 1927), and noise which arises outside the brain, such as a probabilistic relationship between a cue and a reward (Brunswik, 1956). Specifically, our account classifies noise according to when it arises during the information processing that precedes a choice. Encoding noise refers to noise accumulated up to the point at which a stimulus is encoded. As such, encoding noise includes both “external” noise (e.g., a weak correspondence between a retinal image in dim lighting and the object that caused the image) and “internal” noise (e.g., intrinsic stochasticity in sensory transduction). In comparison, integration noise strictly refers to internal noise which arises at later stages of information processing, such as when integrating variable or discordant pieces of information within a limited-capacity system. Under our account, any task that requires the combination of multiple pieces of evidence will be subject to integration noise, and the amount of integration noise will scale with the variability of the different pieces of information that must be combined. Choices may of course be affected by other types of noise than those considered here. For example, cognitive decisions may involve memories, sometimes distant in the past, and risk and ambiguity (Bach & Dolan, 2012; Payzan-LeNestour & Bossaerts, 2011).

Many psychophysical tasks confound encoding and integration noise. For instance, in a random dot-motion task, increasing motion coherence simultaneously increases encoding noise (as instantaneous evidence is less indicative of the correct category of motion) and integration noise (as the variability of evidence across time is higher and thus harder to integrate). Recent work has shown that noisy cognitive inference, related to our notion of integration noise, is a major driver of variability in choices (Drugowitsch, Wyart, Devauchelle, & Koechlin, 2016). Similarly, it has been shown that for complex inference problems, a mismatch between an agent’s internal model of a task and the true structure of a task provokes departures from optimality (Beck et al., 2012). Here we extend these findings by introducing noise blindness as an additional driver of suboptimal cognitive inference. Specifically, the variability in choices caused by integration noise, or by imperfect inference, may not systematically bias choices away from the true choice. Blindness to these sources of choice variability, however, predicts systematic overconfidence, which may manifest itself as a lack of sensitivity to base-rate information, for example. In short, suboptimality can arise not only from having the “wrong” model of the task but also from having the “wrong” model of oneself.

We do not know why humans are blind to integration noise. One possibility is that basing decision strategies on all sources of noise would prolong deliberation and thus reduce reward rates, or that recognising one’s own cognitive deficiencies requires a much longer timeframe. However, a well-known cognitive illusion may help understand why blindness to one’s own cognitive deficiencies may not be catastrophic: even though failures to detect salient visual change suggests that cognitive processing is highly limited (Simons & Levin, 1997), humans enjoy rich, vivid visual experiences of cluttered natural scenes. Human information processing is sharply limited by capacity, but as agents we may not be fully aware of the extent of these limitation.

## Acknowledgments

This work was supported by a Wellcome 4-year-PhD grant to S.H.C. (0099741/Z/12/Z) and an ERC starter grant to C.S. (281628). The Wellcome Centre for Human Neuroimaging is supported by core funding from Wellcome (203147/Z/16/Z).

## Author contributions

S.H.C., D.B., T.E. and C.S. conceived the study. S.H.C., D.B., J.D. and C.S. designed the experiments. S.H.C. programmed the experiments. S.H.C, D.B. and J.D. performed the experiments. S.H.C., D.B. and R.M. developed the models. S.H.C. and D.B. analysed the data and performed the simulations. S.H.C., D.B., J.D., T.E. and C.S. interpreted the results. S.H.C. drafted the manuscript. S.H.C, D.B. and C.S. wrote the manuscript.

## Methods

### Participants

One hundred and five healthy human participants with normal or corrected-to-normal vision were recruited to participate in six experiments (72 females, 8 left-handed, mean age ± SD: 25.02 ± 4.25; Exp1: *n* = 20; Exp2: *n* = 20; Exp3: *n* = 20; Exp4: *n* = 24; Exp5: *n* = 24; Exp6: *n* = 20). Participants were reimbursed for their time and could earn an additional performance-based bonus (see below). All participants provided written informed consent. The experiments were conducted in accordance with local ethical guidelines.

### Experimental paradigm

All six experiments were based on the same psychophysical task. On each trial, participants had to judge whether the average orientation of a circular array of gratings (Gabor patches; see **Fig. 1**) was tilted clockwise (CW) or counter-clockwise (CCW) relative to horizontal. The average orientation of the gratings in each trial was randomly selected from a mixture of two Gaussian distributions (centred at 3° either side of the horizontal axis, respectively, and with 8° of standard deviation). We manipulated encoding noise and integration noise by varying two features of the array in a factorial way manner: the root mean square contrast (*rmc*) of the individual gratings, which affects the difficulty of encoding the stimulus array, and the variability of the orientations of the individual gratings (*std*), which affects the difficulty of integrating orientations across the stimulus array. The number of contrast and variability conditions varied between experiments: in Experiments 1-3, three contrast levels (*rmc* = {0, .16, .6}) and three variability levels (*std* = {0°, 4°, 10°}); in Experiments 4-6, two contrast levels (*rmc* = {.15, .6}) and two variability levels (*std* = {0°, 10°}). The stimulus array was presented for 150 ms and was followed by a 3000 ms choice period. Participants indicated their choice by pressing the right (CW) or the left (CCW) arrow-key on a QWERTY keyboard. They received feedback about choice accuracy, before continuing to the next trial. If no response was registered within the choice period, the word “LATE” appeared at the centre of the screen, and the next trial was started. Experiments 1, 2 and 3 consisted of 1296 trials, divided into 36 blocks of 36 trials each. Experiments 4, 5 and 6 consisted of 1200 trials, divided into 32 blocks of 40 trials each.

In Experiments 1 and 2, participants were presented with a cue to the prior probability of each stimulus category. The cue was presented 700 ms before the onset of the stimulus array and remained on the screen until a response was registered. An “N” indicated that the two stimulus categories were equally likely, an “R” indicated a 75% probability of a CW stimulus and an “L” indicated a 75% probability of a CCW stimulus. Half of the blocks contained neutral trials (“N”) and the other half contained biased trials (“R” or “L”). The blocks were randomised across an experiment. In Experiment 2, after having made a choice, participants were required to indicate the probability that the choice is correct by moving a sliding marker along a scale (50% to 100% in increments of 1%). In Experiment 3, on half of the blocks, participants could opt out of making a choice and receive the same reward as for a correct choice with a 75% probability. There was no prior cue. In Experiment 4, after having made a choice, participants had to categorize (high vs. low) either the contrast or the variability of the stimulus array. Participants received trial-by-trial feedback about the categorisation judgment. The judgment types were counterbalanced across trials. In Experiment 5, for each block of trials, we fixed the contrast or the variability level while varying the other feature. In Experiment 6, on half of the blocks, the stimulus array consisted of eight gratings and, on the other half of blocks, the stimulus array consisted of four gratings. Further experimental details are provided in the Supplementary Information.

### Statistical analyses

All statistics are reported at the group level. We performed analyses of variance (ANOVAs) with participants as a random variable to test the effects of contrast and variability on choice accuracy, response times, cue usage, confidence (Exp2) and opt-in behaviour (Exp3). We performed most analyses of choice accuracy and confidence using neutral trials; analyses of cue usage were naturally based on biased trials. We used multiple linear regression and multiple logistic regression to isolate the effect of variability on confidence and opt-in responses, respectively. For the analyses in **Fig. 5A**, seven participants were excluded because of excessive opt-out responses, but result were almost identical when including them. All p-values lower than .001 are reported as “p < .001”, p-values higher or equal than .001 but lower than .01 are reported as “p < .01”, p-values higher or equal to .01 but lower than .05 are reported as “p < .05”. All p-values greater or equal to .05 are reported as higher than the closest lower decimal (e.g., a p-value of .175 would be reported as “p > 0.1”), with exception of p-values between .05 and . 1 which are reported as “p > .05”. The degrees of freedom for the ANOVAs are specified using non-integer numbers when a Greenhouse-Geisser correction has been used to correct for violations of the sphericity assumption.

### Computational modelling

We first describe the *omniscient* model who takes into account encoding and integration noise and can identify which condition a trial is drawn from (i.e. assigns a probability of 1 to the current condition on a given trial). We then describe the *variability-mixer* model, who takes into account integration noise but cannot distinguish the variability conditions (i.e. assigns equal probability to all variability conditions on a given trial), and the *noise-blind* model, who entirely neglects integration noise. For completeness, we ran six additional models which varied an agent’s awareness of encoding noise and/or ability to discriminate contrast conditions. We only discuss these models in the Supplementary Information as they had no support in the empirical data.

We modelled – regardless of the model – an agent’s noisy estimate, *x*, of the true average orientation, *μ*, as a random sample from a Gaussian distribution with mean *μ* and variance *σ*^2^:

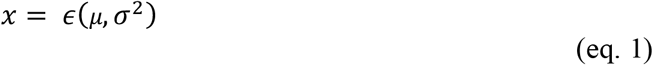

where *σ* is the agent’s total level of noise (encoding plus integration noise) in an experimental condition (see below for noise estimation).

We assumed that an omniscient agent’s internal model has, for each condition, a unique pair of category-conditioned probability density functions (PDFs) over sensory data, which reflect the total level of noise and the true probability distribution over average orientations (see **Fig. 4C** for an example). As such, an omniscient agent would have six pairs of PDFs in Experiments 1-3 and four pairs of PDFs in Experiments 4-6. An omniscient agent uses the relevant pair of PDFs to compute the probability of the sensory data given a CW and a CCW category:

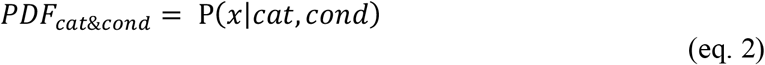

where *cat* is the category and *cond* is the condition. We constructed the PDFs by convolving the true probability distribution over average orientations with a zero-centred Gaussian distribution with variance *σ*^2^ depending on a participant’s total noise in a condition. Note that the construction of these PDFs is specific to the model in question (see construction of “non-omniscient” PDFs below) and is the only source of variation in model predictions about choice and confidence.

We assumed that an agent – regardless of the model – would compute the probability of each category using Bayes’ theorem:

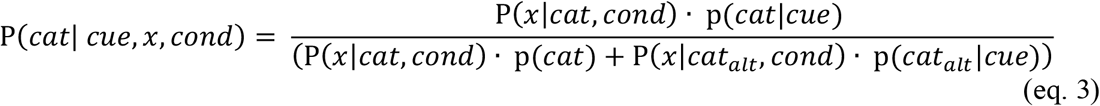

where P(*x*|*cat, cond*) is computed using the relevant PDFs and p(*cat*) is the prior probability of the category in question as indicated by the prior cue. If the category in question is CW, then the alternative category, *cat_alt_* is CCW, and vice versa. On neutral trials, the prior probability of each category is 50%. On biased trials, the prior probability of one category is 75% and the prior probability of the other category is 25%. The computation detailed in eq. 3 can be thought of as scaling the relevant PDFs by the prior probability of the respective category (see **Fig. 4D** for an example).

Finally, we assumed that an agent – regardless of the model – makes a decision, *d*, by selecting the category with higher posterior support and computes confidence in this decision as:

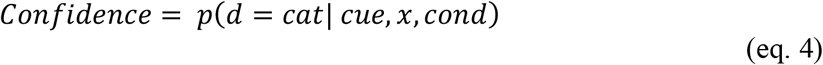

which in our task is directly given by the posterior probability of the chosen category.

Because an omniscient agent takes into account encoding and integration noise and knows which experimental condition a trial is drawn from, she will (i) be appropriately influenced by the prior cue, (ii) accurately estimate the probability of having made a correct choice, and (iii) opt out of trials when she believes that she is less than 75% likely to be correct. We now describe two models which relax the “omniscient” assumptions.

We first consider a *variability-mixer* agent who is sensitive to integration noise but cannot distinguish the different variability conditions. Therefore, when estimating the probability of the sensory data given a CW and a CCW category, the variability-mixer marginalizes its estimate over all possible variability conditions (equivalent to an omniscient agent whose PDFs have been mixed across variability conditions). As a result, when orientation variability is low, the PDFs are more overlapping than for the omniscient model. Conversely, when orientation variability is high, the PDFs are less overlapping than for the omniscient model. For these reasons, a variability-mixer model would display a mixture of under- and overconfidence.

Finally, we consider a *noise-blind* agent who is entirely unaware of integration noise. Like in the case of the variability-mixer model, a noise-blind agent only has a pair of PDFs for each contrast level but, unlike in the case of a variability-mixer model, these PDFs only take into account encoding noise. As a result, when orientation variability is non-zero, the PDFs are less overlapping than under either of the two other models (**Fig. 4E**) and a noise-blind agent would therefore tend to hold stronger posterior beliefs (i.e. steeper curves for Fig.4F). Such stronger posterior beliefs will lead a noise-blind agent to (i) be less influenced by the prior cue than needed, (ii) overestimate the probability of having made a correct choice, and (iii) not opt out of trials when being less than 75% likely to be correct.

We note that the models make the same predictions about choice on neutral trials but are distinguishable when focusing on (i) biased trials and (ii) confidence and opt-in behaviour on both neutral and biased trials. Our modelling approach allowed us to calculate a choice probability for each trial under a given model. For model analyses requiring a categorical choice (e.g., logistic regression), we sampled choices according to these choice probabilities.

### Noise estimation

We assumed that each experimental condition was affected by Gaussian noise with a specific standard deviation, *σ_cond_*. We assumed that encoding noise depends upon the contrast of the array and that integration noise is proportional to the variability of orientations in the array. We estimated the total level of noise for each condition using four free parameters (three for Experiments 4-6). Two parameters characterised the level of encoding noise for each contrast level: one for low contrast (nC_low_) and one for high contrast (nC_high_). The other two parameters (one for Experiments 4-6) characterised the level of integration noise for each variability level: one for medium variability (nV_med_, only for Experiments 1-3) and one for high variability (nV_high_). For a given condition, the total level of noise (the standard deviation of the Gaussian noise distribution), *σ_cond_*, is thus given by:

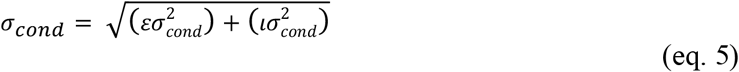

where *εσ_cond_* and *ισ_cond_* specify the contribution of encoding noise and integration noise, respectively. For instance, for the low-contrast, high-variability condition would be given by substituting nC_low_ for *εσ_cond_* and nV_high_ for *iσ_cond_*.

We fitted the four noise estimators for each participant by maximizing the likelihood of the participant’s choice using neutral trials only (we used a genetic algorithm with a population size of 100 individuals and a maximum generation time of 1000 generations). We note that, because of our factorial design, we could separate the two sources of noise. We used the fitted parameters for each participant to construct the model PDFs described above. We stress that the noise estimation use choices on neutral trials only and that the model predictions pertain to independent features of the data: (i) confidence on neutral trial choices, (ii) choices (and choice probabilities) on biased trials, and (iii) probability of opting out.

The mean ± SEM of the best fitting values for the four noise parameters (nC_low_, nC_high_, nV_med_ and nV_high_) in units of degrees were: 10.10 ±1.51, 3.31 ± 0.39, 3.0 ± 0.78 and 6.8 ± 1.0, respectively. Following equation 5, the estimated total amounts of noise fitted for the three key conditions (baseline, low-c and high-v) were therefore: 3.31 ± 0.39, 10.1 ± 1.51 and 8.0 ± 1.0, respectively. There was a significant difference between the values for the baseline condition and those for the other two conditions (both p-values < 0.001), but no significant difference between the low-c and high-v conditions (p-value > 0.16).

### Psychometric fits

We fitted psychometric curves to the average proportion of clockwise choices using a four-parameter logistic function:

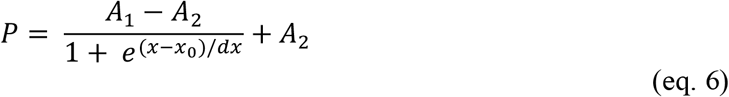

where *P* is the proportion of CW choices, *A*_1_ is the right asymptote, *A*_2_ is the left asymptote, *x*_0_ is the inflection point and 1/*dx* is the steepness, and *x* is the average stimulus orientation at which the proportion of CW choices is evaluated. We computed the proportion of clockwise choices within average-orientation bins (i.e. six quantiles over the average orientation relative to horizontal). The psychometric curves shown in **Fig. 2B** are only used for illustration.

### Bias index

We used Signal Detection Theory (Macmillan & Creelman, 2004; Stanislaw & Todorov, 1999) to calculate the decision criteria, *c*, separately for trials on which the prior cue favoured CW and trials on which the prior favoured CCW. The decision criterion provides a signed estimate of the degree to which the prior cue biases a participants’ choices independently of their sensitivity to average orientation. We computed the criterion as, *c* = −0.5[*Φ*^−1^(*HR*) + *Φ*^−1^(*FAR*)], where *Φ*^−1^ represents the inverse of the normal cumulative density function, and HR and FAR represent the hit rate (i.e. the proportion of CW trials where participants responded CW) and false alarm rate (i.e. the proportion of CCW trials where participants responded CW), respectively. We then used the difference between *c* when cued CW (*c*_CW_) and *c* when cued CCW (*c*_CCW_) as our measure of cue usage: bias index = *c*_CW_-*c*_CCW_. Higher values indicate greater cue usage. We computed a bias index for each participant and each experimental condition.

## Supplementary Information

### Experimental details

In all experiments, participants had to judge the average orientation of an array of gratings as clockwise (CW) or counter-clockwise (CCW) from horizontal. We first describe trial events, trial timings and stimulus construction for Experiment 1 and then explain the additional steps taken for Experiments 2-6.

In Experiment 1, a fixation dot first appeared at the centre of the screen for 300 ms to announce the start of a trial. The fixation dot was replaced by a cue which appeared 700 ms before the stimulus array and which remained on the screen until a response was registered. The cue determined the prior probability of each stimulus category (“L”: prior probability of CCW is 75%; “N”: CCW and CW equally likely; “R”: prior probability of CW is 75%). The stimulus array was shown for 150 ms and was followed by an up-to 3000 ms long response window. Participants responded by pressing the left (CCW) or right (CW) arrow-key on a QWERTY keyboard using their right hand. Categorical feedback about choice accuracy (“CORRECT” or “WRONG”) appeared once a response had been registered and remained on the screen for 500 ms, before the onset of the next trial. If no response was registered within the response period, the word “LATE” appeared at the centre of the screen for 3000 ms, and the next trial was automatically started.

The stimulus was composed of eight gratings displayed within a circular array. We manipulated two features of the stimulus array in a factorial manner: the contrast of the gratings and the variability of the gratings’ orientation.

The centre of each grating was located at a distance of ~4.3 degrees of visual angle (400 pixels) from the centre of the screen. Each grating was a Gabor patch constructed using the following parameter values: diameter of ~1.07 degrees of visual angle (100 pixels); spatial frequency of ~5 cycles per degree of visual angle (0.05 cycles per pixel); random phase. All gratings had the same root mean square contrast (rmc, henceforth contrast). The contrast of a trial was either 0 (no signal), .15 (low contrast) or .60 (high contrast). The latter two contrast levels may not affect orientation discrimination on their own (Mareschal & Shapley, 2004). However, we added low-level random noise to the gratings (Wyart, Nobre, & Summerfield, 2012). For each grating, we convolved a unique patch of white noise with a two-dimensional zero-mean Gaussian with a standard deviation of ~0.21 degrees of visual angle (20 pixels). The amplitude of the noise was 10% of the maximum possible. We then added the noise to the grating. Finally, we convolved the grating with a 2-dimensional Gaussian envelope peaking at the centre of the grating and decaying with a standard deviation of ~0.21 degrees of visual angle (20 pixels).

The average orientation of gratings on a trial (henceforth trial mean) was randomly drawn from a Gaussian distribution with a mean of −/+ 3° and a standard deviation of 8°. The variability in the orientations of gratings on a trial (henceforth variability) was randomly drawn from a Gaussian distribution with a mean 0° and a standard deviation of either 0° (zero variability), 4° (medium variability) or 10° (high variability). To ensure the trial mean remained unchanged after the variability manipulation, we subtracted the mean deviation from 0 from the gratings’ orientations. Together, these steps allowed us to independently manipulate trial mean, contrast and variability. We emphasise that feedback was determined by the average orientation of the gratings presented and not by the distribution from which they were drawn.

The experiment consisted of 1296 trials, distributed into 36 blocks of 36 trials each. On half of the blocks, the prior cue was “N” (neutral trials). On the other half of blocks, the cue varied between “L” or “R” in a trial-by-trial manner (biased trials). Block order was randomised across an experiment and across participants.

In Experiment 2, we introduced an explicit measure of confidence. Participants indicated their choice by pressing “Z” (CCW) or “X” (CW) using their left hand. After having made a choice, participants had to indicate the probability that the choice is correct. Participants indicated their confidence by sliding a marker along a vertical scale (50% to 100% in increments of 1%) using a standard computer mouse with their right hand. The probability associated with the marker’s current position was updated in real-time and shown at the centre of the screen. Participants confirmed their response by clicking the left button of the mouse. There was no time limit for the confidence judgment. Feedback about choice accuracy appeared 300 ms after a response had been confirmed. Trial numbers, block types and structure were the same as for Experiment 1.

In Experiment 3, we introduced an implicit measure of confidence. On half of the blocks, participants could choose to opt out of making a choice and receive the same reward as a correct choice with a 75% probability. To remind participants about the choice options on a trial, the words “LEFT” (CCW) and “RIGHT” (CW) appeared to the left and the right of the fixation cross after the stimulus disappeared and, when the opt-out option was available, the words “OPT OUT” appeared below the fixation cross. The opt-out option was selected by pressing the downwards arrow key. For feedback, “SUCCESS” was shown after a correct choice and a rewarded opt-out response, whereas “FAILURE” was shown after an incorrect choice and an unrewarded opt-out response. The experiment consisted of 1296 trials, distributed into 36 blocks of 36 trials each. On half of the blocks, the opt-out option was not available. On the other half of the blocks, the opt-out option was available. Block order was randomised across an experiment and across participants. There was no prior cue.

In Experiment 4, we asked participants to categorise either the contrast (rmc = {.15, .60}) or the variability (std = {0°, 10°}) of the stimulus array. In particular, after having made a choice (by pressing the “X” and “Z” buttons using their left hand, with the chosen category highlighted in bold), participants were then required to judge whether the contrast of the stimulus array was high or low or whether the variability of the stimulus array was high or low. The relevant stimulus dimension for the second judgment (indicating by displaying “CONTRAST” or “VARIABILITY at the centre of the screen), was determined randomly and was only revealed after an orientation discrimination had been made. Participants made the second judgment by pressing the left (low) or the right (high) arrow key (the options “LOW” and “HIGH” appeared equidistantly to the left and the right of the fixation point). Once participants had made their response, they received feedback about the accuracy of each judgment, indicated by changing the colours of the selected options to red (incorrect) or green (correct). The experiment consisted of 1200 trials, distributed into 32 blocks of 40 trials each. On half of the blocks, the prior cue was “N” (neutral trials). On the other half of blocks, the cue varied between “L” or “R” in a trial-by-trial manner (biased trials). Block order was randomised across an experiment and across participants.

In Experiment 5, we fixed either contrast or variability across blocks of trials. Specifically, within a block, one dimension was fixed (low or high), while the other dimension varied randomly (low or high). For instance, in one condition, contrast would be fixed at low while variability varied between high and low across the trials within the block. As in Experiment 4, there were only two levels of contrast and two levels of variability. There were thus eight blocks for each trial type. On half of the blocks for a trial type, the prior cue was “N” (neutral trials). On the other half of blocks for a trial type, the prior cue varied between “L” or “R” in a trial-by-trial manner (biased trials). The experiment consisted of 1200 trials, distributed into 32 blocks of 40 trials each. Block order was randomised across an experiment and across participants.

In Experiment 6, we varied the set-size of the stimulus array. In particular, the stimulus array was composed of either four or eight gratings. As in Experiments 4 and 5, there were only two levels of contrast and two levels of variability. For arrays with only four gratings, the location of the gratings was fixed within a block but randomised across blocks, by sampling a random set of four contiguous locations from the full array with eight gratings. The set-size was varied in a trial-by-trial manner. Half of the trials had a set-size of four gratings and the other half of trials had a set-size of eight gratings. We scaled the variance of the distribution of orientation variabilities for the four-item set size to ensure that the observed standard deviation of orientations within a set-size was equated for the four-item and eight-item case. Without this step, the observed variance of the four-item set size would be systematically lower than for the eight-item set-size. The experiment consisted of 1200 trials, distributed into 32 blocks of 40 trials each. On half of the blocks, the prior cue was “N” (neutral trials). On the other half of blocks, the cue varied between “L” or “R” in a trial-by-trial manner (biased trials). Block order was randomised across an experiment and across participants.

All participants were reimbursed for their participation and had the opportunity to earn an additional performance-based bonus. In all experiments except Experiment 2, participants received a flat rate of £10 and could earn an additional £1 for every 2% increase in choice accuracy relative to 60%. In Experiment 2, participants received a flat rate of £5 and could earn an additional bonus depending on the accuracy of their confidence judgments. We submitted participant’ responses to a strictly proper scoring rule under which it was in participants’ best interest to make as many correct decisions as possible and to estimate the probability that their choice is correct as accurately as possible (Sonnemans & Theo Offerman, 2001). The average bonus accrued was ~£12.

Participants received a 10-minute introduction to their corresponding task, including the stimulus, sources of choice difficulty, prior cue and prior probabilities, response contingencies, and the rules behind the performance-based bonus. Participants also completed a short practice session (two blocks) of the task before starting the experiment proper.

### Category-conditioned probability density functions for an omniscient agent

We assumed that an omniscient agent’s internal model has, for each experimental condition, a unique pair of category-conditioned probability density functions (PDFs) over sensory data. An example of a full set of PDFs are shown in **Fig. S1**. Note that the PDFs look more skewed in conditions with low noise (top-left in **Fig. S1**) as they will more closely resemble the true distribution of average orientations (**Fig. 4A**).

**Fig. S1.**
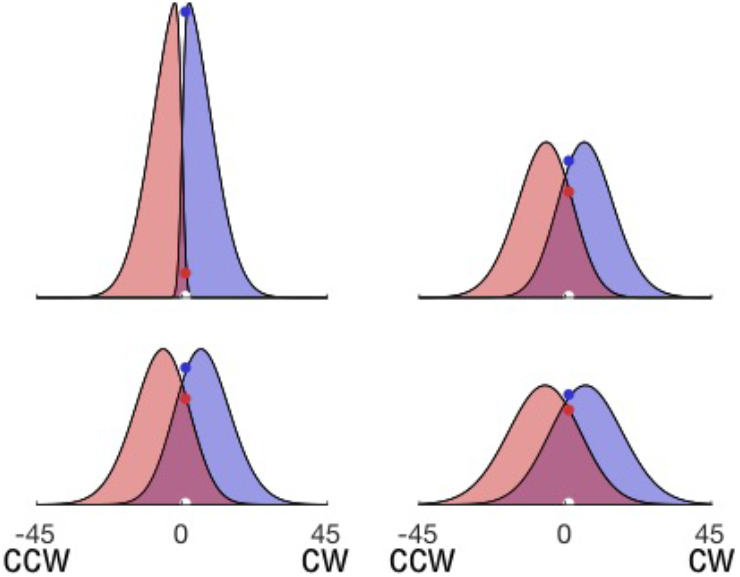
Category-conditioned probability density functions. Four pairs of PDFs, one for each of the four conditions in Experiment 4-6 (but six pairs of PDFs for Experiments 1-3). Top-left: high-contrast and low-variability trials (lowest noise). Top-right: high-contrast and high-variability trials (intermediate noise). Bottom-left: low-contrast and low-variability trials (intermediate noise). Bottom-right: low-contrast and high-variability trials (highest noise). The white dots on the x-axes denote an agent’s current sensory data (same across panels). The blue and red dots indicate the probability density that the sensory data came from a CW or a CCW category, respectively.

### Comparison of computational models

All models considered share the same generative (true) model of how sensory data is generated but differ in their internal model of this process. In particular, they differed with respect to (i) an agent’s ability to identify which condition a trial is drawn from and (ii) an agent’s sensitivity to the different sources of noise in play. These differences give rise to different inferences and thereby responses. We compared a total of nine candidate models of our data. In Table S1, we provide the model names (column 1), quantitative comparison against the noise-blind model which was the best-fitting model (column 2), and details about model assumptions (columns 3-9). Note that the number of unique pairs of category-conditioned PDFs depends on the experiment (the first number is for Exp1-3; the number in brackets is for Exp4-6). For completeness, we considered models which intuitively seemed unlikely (e.g., the Full Mixer model who cannot tell at all discriminate experimental conditions), or are not grounded on optimality (e.g. the models that operate with the average noise across conditions).

**Table S1.** Model assumptions and model comparison. Model comparison was based on the difference in average BIC across participants (Exp1-3) relative to the noise-blind model.

| Model Name | Average BIC difference with best model | Number of unique pairs of PDFs | Is blind to Encoding Noise? | Knows the contrast condition of each trial? | Is blind to Integration Noise? | Knows the variability condition of each trial? | Operates with the average Encoding Noise? | Operates with the average Integration Noise? |
| --- | --- | --- | --- | --- | --- | --- | --- | --- |
| Omniscient | -32.98 | 6 (4) | no | yes | no | yes | no | no |
| Noise Blind | 0.00 | 2 (2) | no | yes | yes | irrelevant | no | no |
| Variability Mixer | -20.41 | 2 (2) | no | yes | no | no | no | no |
| Contrast Mixer | -60.39 | 3 (2) | no | no | no | yes | no | no |
| Full Mixer | -58.57 | 1 (1) | no | no | no | no | no | no |
| Average Variability | -47.28 | 2 (2) | no | yes | no | no | no | yes |
| Average Contrast | -116.93 | 3 (2) | no | no | no | yes | yes | no |
| Full Average | -134.76 | 1 (1) | no | no | no | no | yes | yes |
| Contrast Blind | -9.79 | 3 (2) | yes | irrelevant | no | yes | no | no |

### Overconfidence in choices

A main difference between the variability-mixer and the noise-blind models is the predicted pattern of *overconfidence* (i.e. mean confidence minus mean accuracy) across the key conditions of our factorial design. The variability-mixer model predicts a hard-easy effect, with overconfidence for the high-variability condition and underconfidence for the baseline condition. By comparison, while the noise-blind model also predicts overconfidence for the high-variability condition, it predicts good calibration for the baseline condition. Indeed, as expected under the noise-blind model, participants were overconfident in the high-variability condition but well-calibrated in the baseline condition (Fig S2).

**Fig. S2.**
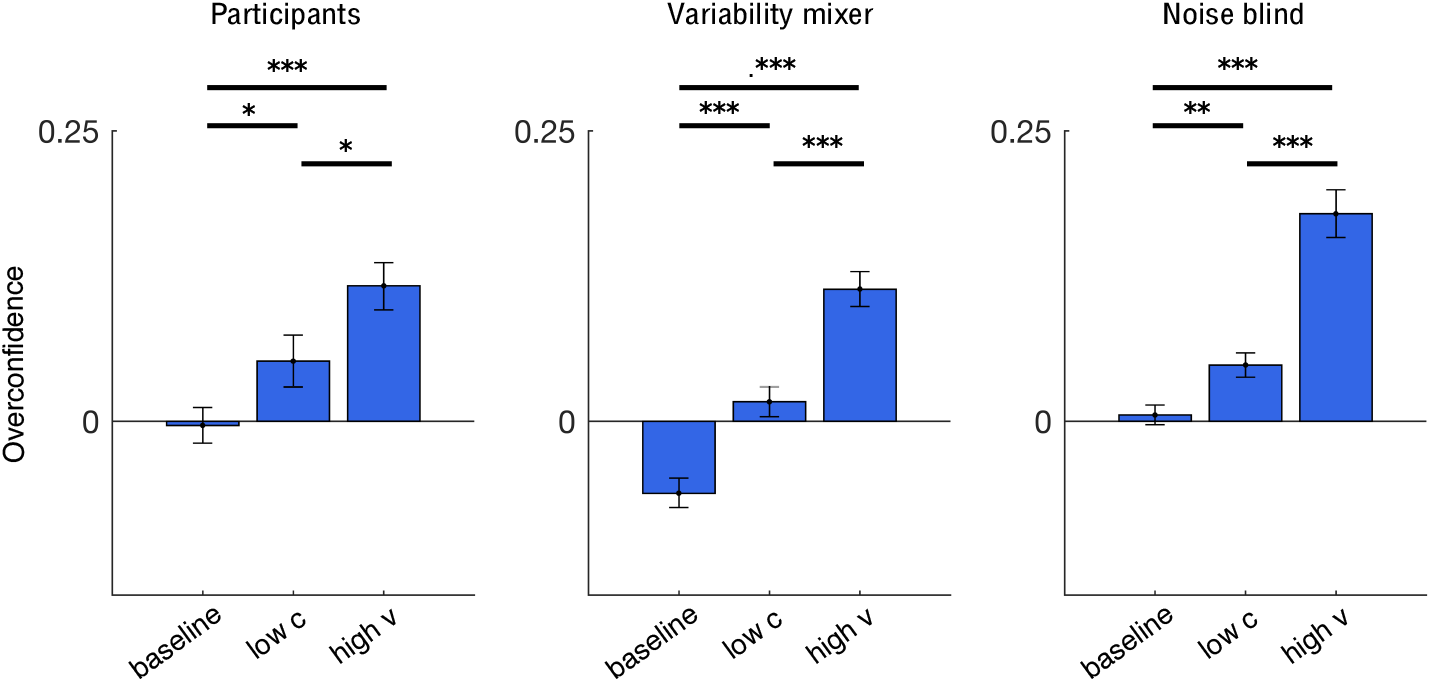
Noise-blind model accounts for empirical pattern of overconfidence. Participants (left) are well-calibrated in the baseline condition but overconfident in the other conditions. The variability-mixer model (middle) shows under-confidence in the baseline condition but overconfidence in the high-variability condition. The noise-blind model shows good calibration in the baseline condition and overconfidence in the high-variability condition. Near-zero values indicate good calibration and non-zero values indicate bad calibration. Negative values indicate under-confidence and positive values indicate overconfidence. Data is from neutral trials of Experiment 2 and represented as group men ± SEM.

### Sequential sampling model

To test the sequential sampling account of suboptimal behaviour proposed by Zylberberg and colleagues (Zylberberg et al., 2016), we fitted a Drift Diffusion Model (DDM) to the data from Experiments 1-2. The DDM models two-choice decision-making as a process of accumulating noisy evidence over time with a certain speed, or drift-rate, until one of two choice thresholds is crossed and the associated response is executed. We assumed that the choice thresholds were fixed across experimental conditions as in the study by Zylberberg and colleagues. In addition, we assumed that lower contrast led to a lower mean of the drift-rate and that higher variability led to higher variance of the drift-rate. The base drift-rate was proportional to the absolute difference between the average orientation and horizontal. We implemented these mechanisms using three parameters. The first parameter depends on the contrast level and scales the drift-rate. The second parameter specifies the baseline variance of the drift-rate. Finally, the third parameter depends on the variability level and scales the baseline variance of the drift-rate. To find the best parameters for each participant, we minimized the sum of squared errors between empirical and predicted choice accuracy across experimental conditions (we used a genetic algorithm with a population size of 1000 individuals and a maximum generation time of 1000 generations). Comparison between empirical data and model predictions are shown in **Fig. S3** using a similar analyses as Zylberberg and colleagues (2016). In short, the model can predict the observed choice accuracy for the different conditions (**Fig. S3A**), but it predicts a pattern of response times with respect to stimulus variability opposite to what we observed (**Fig. S3B**). As a sanity check, we show that higher evidence strength (i.e. absolute deviation of the average orientation from the category boundary) indeed increases choice accuracy and fastens response times (**Fig. S3C-F**).

**Fig. S3.**
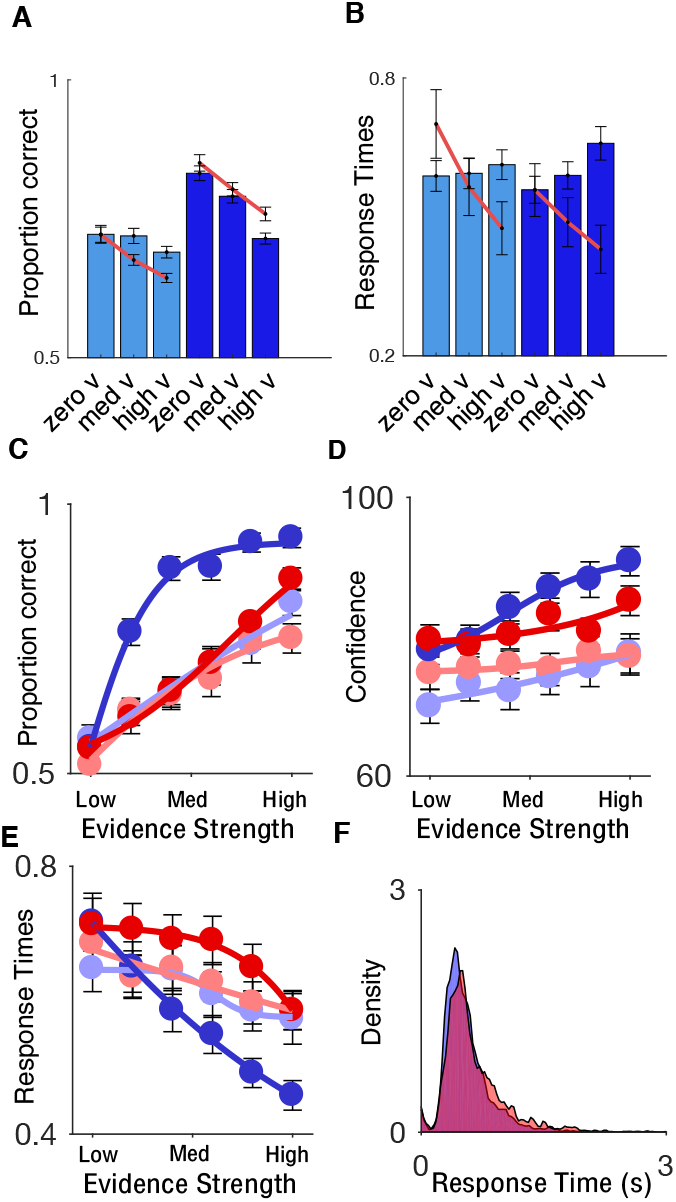
Common choice threshold in a sequential sampling model cannot explain our data. (**A**) Choice accuracy for participants (blue bars) and DDMs (red lines) is lower when contrast is low (compare pale blue and dark blue bars) and when variability is high (negative slopes as condition changes from zero-v to high-v). (**B**) Response times for participants and DDMs show opposite effects for increases in variability (positive slopes for participants but negative ones for DDMs). (**C**) Participants’ choice accuracy for different levels of evidence strength (blue: low variability, red: high variability; faint colours: low contrast; dark colours: high contrast). Note that the two critical conditions, high-variability and high-contrast trials (dark red curve) and low-contrast and zero-variability trials (faint blue curve), have similar slopes. (**D**) Participants’ confidence for different levels of evidence strength (same colour scheme as in panel c). (**E**) Participants’ response times for different levels of evidence strength (same colour scheme as in panel c). (**F**) Collapsing response times across participants for high-contrast and zero-variability trials (blue) and high-contrast and high-variability trials (red) demonstrate that high variability is associated with slower response times (i.e. red distribution has a longer tail). Data is from neutral trials of Experiments 1-2 and represented as group men ± SEM. For panels C-E, evidence strength is divided into quantile bins of roughly one degree of width starting at zero.

### Subsampling

In our experiments, we observed a consistent decrease in performance for trials with high stimulus variability. We attributed this decrease to integration noise – an increased difficulty for integrating variable or disparate pieces of information – which can explain both decreased choice accuracy and increased response times for high-variability stimuli. An alternative explanation for the decrease in accuracy for high-variability stimuli is, however, that participants only based their judgments on a subset of the items in a stimulus array. Under this subsampling account, the decrease in accuracy is due to a larger mismatch between the actual average orientation of the full array and the average orientation of the sampled subset. Here we describe why subsampling is an unlikely explanation for the decrease in accuracy for high variability trials, and why subsampling cannot explain noise blindness.

First, we found no effect of set-size on accuracy in Experiment 6 (**Fig. S4A**) which was designed to show a difference in performance between set-sizes if participants were indeed subsampling. More specifically, in this experiment, the distribution of average orientations was the same for both set-sizes. Therefore, if participants sampled all items, then there would be no *mismatch* and thus no difference in performance between the two set-sizes, and the decrease in performance between the *baseline* and *high-v* conditions cannot be explained by subsampling. If, on the other hand, subsampling did occur, it would have a bigger effect on performance on trials where the stimulus arrays are composed of eight-item than on trials with four items. For instance, if participants could sample four items, then there would be no difference in performance between the *baseline* and the *high-v* conditions for four-item arrays (because there would be no mismatch), but there would be a difference for eight-item arrays (because half of the items would have been ignored). Experimentally, we found that the decrease in performance between the *baseline* and the *high-v* conditions was consistent across set-sizes and comparable to that found in the previous experiments (**Fig. S4A**). We note that another prediction for Experiment 6 is that accuracy should be higher on eight-item than four-item trials because encoding noise could be averaged out over more items. However, the data does not support this prediction. One possibility is that there is a trade-off between the number of items that are encoded and the quality with which they are encoded (Van den Berg, Shin, Chou, George, & Ma, 2012) – a trade-off which may overshadow the expected boost in performance from averaging out encoding noise.

**Fig. S4.**
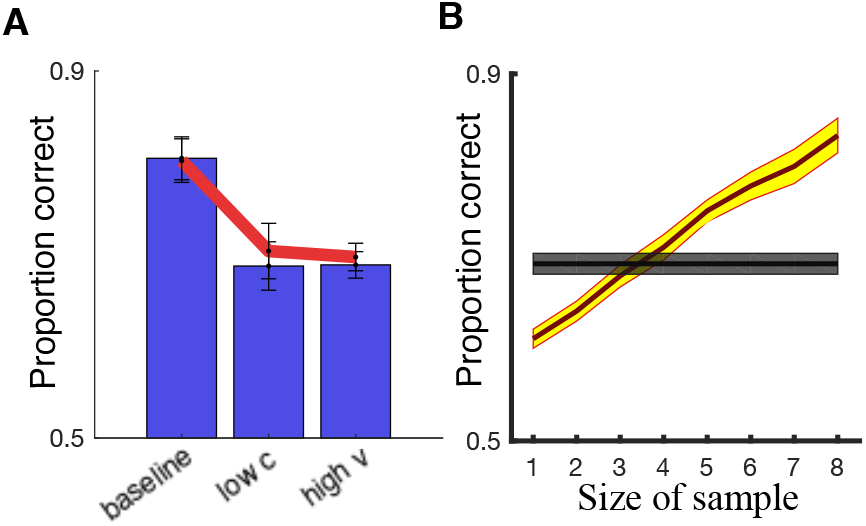
Noise blind and not subsampling. **(A)** Participants in Experiment 6 achieved similar levels of performance when required to integrate four items (blue bars) and eight items (red line). **(B)** Choice accuracy for participants (grey shade) and a subsampling model (yellow shade). The subsampling model, which has no integration noise and therefore perfectly averages the sampled gratings, would need to sample about four gratings before reaching the same level of choice accuracy as participants. (A-B) Data is represented as group mean ± SEM.

Second, we performed a set of simulations for a subsampling agent without integration noise where we varied the number of items sampled (**Fig. S4B**). As such, the simulations were carried out assuming that, for arrays made up of eight gratings, the decrease in accuracy between the baseline and high-v conditions was entirely driven by subsampling. While sampling four-items could in principle explain the decrease in accuracy between the baseline and high-v conditions for eight-item arrays, the same number of sampled items would represent the complete stimulus for arrays of four-item arrays and no expected decrease in performance should be found for high-v compared to baseline trials.

Third, we fitted a subsampling model to participants’ data (only neutral trials) to directly quantify the number of items sampled by each participant (*n* = 60; Exp1-3). The model had three free parameters. The first parameter controls the noise added to each item of the array in the baseline condition. The second parameter controls the extra amount of noise added to each item in trials where the contrast is low (to capture the extent of encoding noise). Finally, the third parameter controls the number of gratings, k, that were sampled from a stimulus array; *k* is an integer value between one (the minimum number of items that can be sampled) and eight (the total amount of items that can be sampled). We fitted the parameters by maximising the likelihood of participants’ choices using a genetic algorithm with a population size of 100 individuals and a maximum generation time of 1000 generations. Note that this subsampler account does not have integration noise and any reduction in accuracy for high-variability stimuli would have to be due to subsampling. Even then, the fitted *k*-parameter was eight for most participants (Table S2).

**Table S2.**
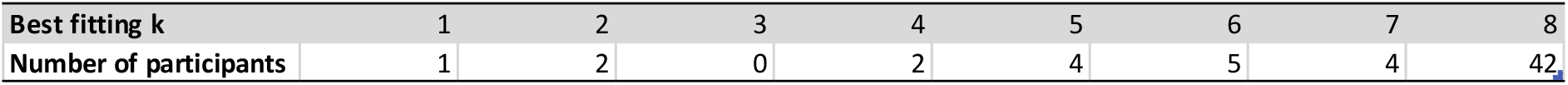
Estimated number of items, *k*, sampled by participants assuming absence of integration noise.

In summary, subsampling is unlikely to explain the observed reduction in accuracy for trials with high stimulus variability, and provides no explanation for the slower responses observed by participants or for the apparent blindness to the performance cost associated with high stimulus variability. We therefore argue that integration noise provides the best account of the reduction in accuracy and the delay in responses observed for trials with high stimulus variability, and that blindness to this noise is the most parsimonious explanation of the overconfidence, lack of usage of the opt-out option and lack of influence of the prior cue observed for trials with high stimulus variability.

### Response times

In many tasks, response times provide experimenters with further information about the computational processes that lead to a decision, with response times varying with the 1) difficulty of a decision as well as 2) the confidence with which it was made.

First, difficult decisions require more deliberation, and responds times therefore tend to be slower for harder stimuli. In our task, response times were indeed slower for the low-c and the high-v conditions compared to the baseline condition (baseline<low-c: *t*(39) = 2.6, *p* < .05; baseline<high-v: *t*(39) = 6.15, *p* < .001; see **Fig. S5**). However, response times were even slower for the high-v compared to the low-c condition (low-c<high-v: *t*(39) = 4.0, *p* < .001), despite equal levels of choice accuracy in the two conditions. Analysis of the full data set (ANOVA) confirmed that response times increased with variability (main effect of variability: *F*(1.5,29.0) = 28.55, *p* < .001), whereas response times did not vary directly with contrast (main effect of contrast: *F*(1,39) = 0.09, *p* > .75), only through an interaction with variability (*F*(1.8,72.7) = 13.06, *p* < .001). Overall, our argument that integration noise results from an increased difficulty for integrating variable or discordant pieces of information is supported by the slower response times observed for conditions with high stimulus variability.

Second, there are two different ways of thinking about the relationship between confidence to response times. On one hand, we – the experimenters – can use response times as a proxy for participants’ confidence. However, this relationship may not be straightforward; quick responses could reflect either rapid guesses or high certainty, and slow responses could reflect thoughtful deliberation or high uncertainty (Pleskac & Busemeyer, 2010). On the other hand, participants may themselves use the time it took them to make a decision as a cue to how likely their decision is to be correct. By recording both response times and confidence judgments, we could investigate the contribution of response times to confidence over and above relevant stimulus features (average orientation, contrast and variability). The incentive-compatible scoring procedure applied to participants’ responses meant that participants, to maximise reward, should make as many correct decisions as possible and estimate the probability that a choice is correct as accurately as possible. As demonstrated by the trial-by-trial analysis of confidence presented in **Fig. 3B**, slower response times were indeed associated with a decrease in confidence (see **Fig. 3B**). In other words, participants utilised response times as a cue to confidence. However, the analysis also shows that, because there were additional influences on confidence (e.g., average orientation and variability). response time is a poor proxy for participants’ confidence.

**Fig. S5.**
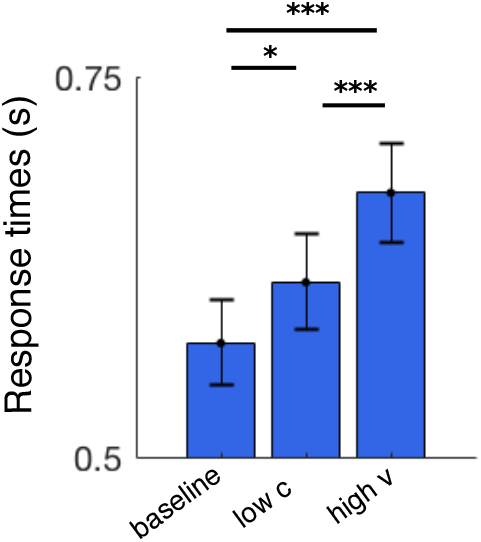
Response times for critical conditions. Response times were fastest for the baseline condition, and slowest for the high-v condition. Data is from neutral trials (Exp1-2) and represented as group mean ± SEM.

### Accuracy gain

In our task, participants had an opportunity to compensate for poor performance when the informative prior cue (Exp1-2) or an opt-out option(Exp4) was available. Under the noise-blindness account, such an accuracy gain on ‘extra-information’ trials compared to neutral trials should be higher for the low-c than the high-v condition, unless participants employed other strategies which allowed them to compensate for the errors associated with stimulus variability (e.g., by deliberating for longer at the expense of slower responses). To test these predictions, we computed the difference in choice accuracy between ‘extra-information’ and neutral trials: *Accuracy_gain_* = *Accuracy_extra_information_* − *Accuracy_neutral_*. In line with our hypothesis, accuracy gains (**Fig. S6**) were higher on high-c than high-v trials for participants and the noise-blind model, but not for the omniscient and the variability-mixer models.

**Fig. S6.**
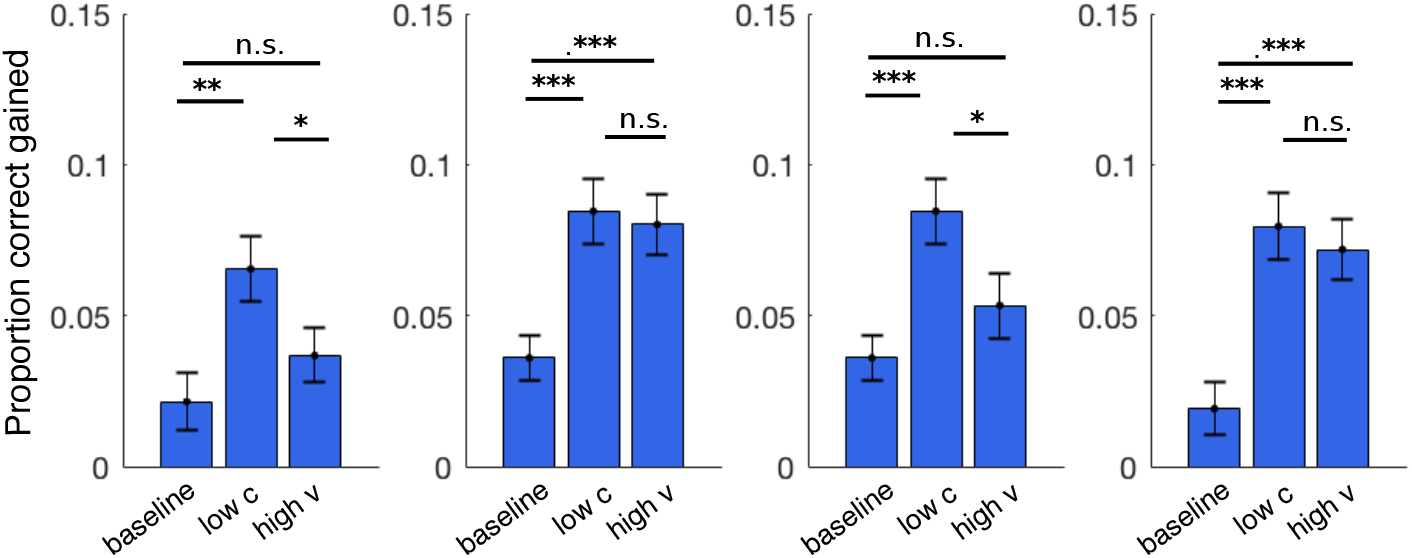
Accuracy gain for critical conditions. Accuracy gain is measured as the difference in choice accuracy between ‘extra-information’ and neutral trials (with positive values meaning higher accuracy for ‘extra-information’ trials). From left to right, the panels show the accuracy gain for participants, the omniscient model, the noise-blind model, and the variability-mixer model, respectively. Data is from Exp1 - 3 and represented as group mean ± SEM.

